# Improving the integration of AI into existing ecological inference workflows

**DOI:** 10.1101/2024.11.27.625677

**Authors:** Amber Cowans, Xavier Lambin, Darragh Hare, Chris Sutherland

## Abstract

1. Artificial Intelligence (AI) has revolutionised the process of identifying species and individuals in audio recordings and camera trap images. However, despite developments in sensor technology, machine learning, and statistical methods, a general AI-assisted data-to-inference pipeline has yet to emerge.
2. We argue that this is, in part, due to a lack of clarity around several decisions in existing workflows, including: the choice of classifier used (e.g., semi- vs. fully automated); how classifier confidence scores are used and interpreted; and the availability and selection of appropriate statistical methods for drawing ecological inferences.
3. Here, we attempt to conceptualise a general workflow associated with automated tools in ecology. We motivate this perspective using our experiences with occupancy modelling using monitoring data collected through passive acoustic monitoring and camera trapping, identifying priority areas for future developments.
4. We offer an accessible guide to support the ecological community in navigating and capitalising on rapid technological and methodological advances. We describe how different error types arise from both sensor-based monitoring and from classifiers themselves; how different error types are handled at each stage of the workflow; and finally, implications and opportunities associated with deciding on methods used at each step of the pipeline.
5. We recommend that “black box” tools like neural network classification algorithms should be embraced in ecology, but widespread uptake requires more formal integration of AI into the existing ecological inference workflows. Like ecological AI more broadly, however, successful development of new data-to-inference pipelines is a multidisciplinary endeavour that requires input from everyone invested in collecting, processing, analysing, and using ecological monitoring data.

## Detection and classification for ecological data

Advances in remote sensing technologies, automated classification methods, and statistical models for observation processes have revolutionised the study of the natural world (Besson et al., 2022). Ecological sensors such as camera traps and autonomous recording units (ARUs) are now routinely deployed to monitor species’ distribution (Chambert, Waddle, et al., 2018), abundance (Augustine et al., 2018; Palencia et al., 2022) and behaviour (Buxton et al., 2018; Caravaggi et al., 2017), and method-specific uncertainties are well-documented (Burton et al., 2015). A growing and widely applied suite of hierarchical models, particularly occupancy models, have been developed to draw ecological inferences from sensor data while accounting for false negative errors: the ubiquitous failure to detect a species or individual when present, and false positives: the less common, but still problematic, erroneous detection of a species or individual when absent (Kéry & Royle, 2020; Royle & Link, 2006). These models typically require discrete representations of monitoring data (binary detections or counts) and focus on correcting for imperfect detection by sensors, i.e., the *detection* process (MacKenzie et al., 2002; Rota et al., 2016), and more recently for species or individual misclassification, i.e., the *classification* process (Chambert, Waddle, et al., 2018; Wright et al., 2020).

Unlike the binary detections generated using manual labelling by humans, machine learning classifiers assign continuous confidence scores (0 < *c* < 1) to a list of potential identities (e.g., species or individuals), where scores closer to 1 are assumed more likely to be correct (Dussert et al., 2024; Kéry & Royle, 2020; Rhinehart et al., 2022). While the probability of detection on a sensor is unaffected, the introduction of AI into existing workflows alters how data are processed and, importantly, how they should be analysed (Figure 1). Most commonly, these confidence scores are converted to binary detections based on arbitrarily defined thresholds and then traditional modelling approaches are applied, assuming the data are truly binary (Fiss et al., 2024; Lonsinger et al., 2023; Whytock et al., 2021). In contrast, recently proposed coupled classification models address classification uncertainty by directly modelling these continuous confidence scores (Kéry & Royle, 2020; Rhinehart et al., 2022).

**Figure 1.**
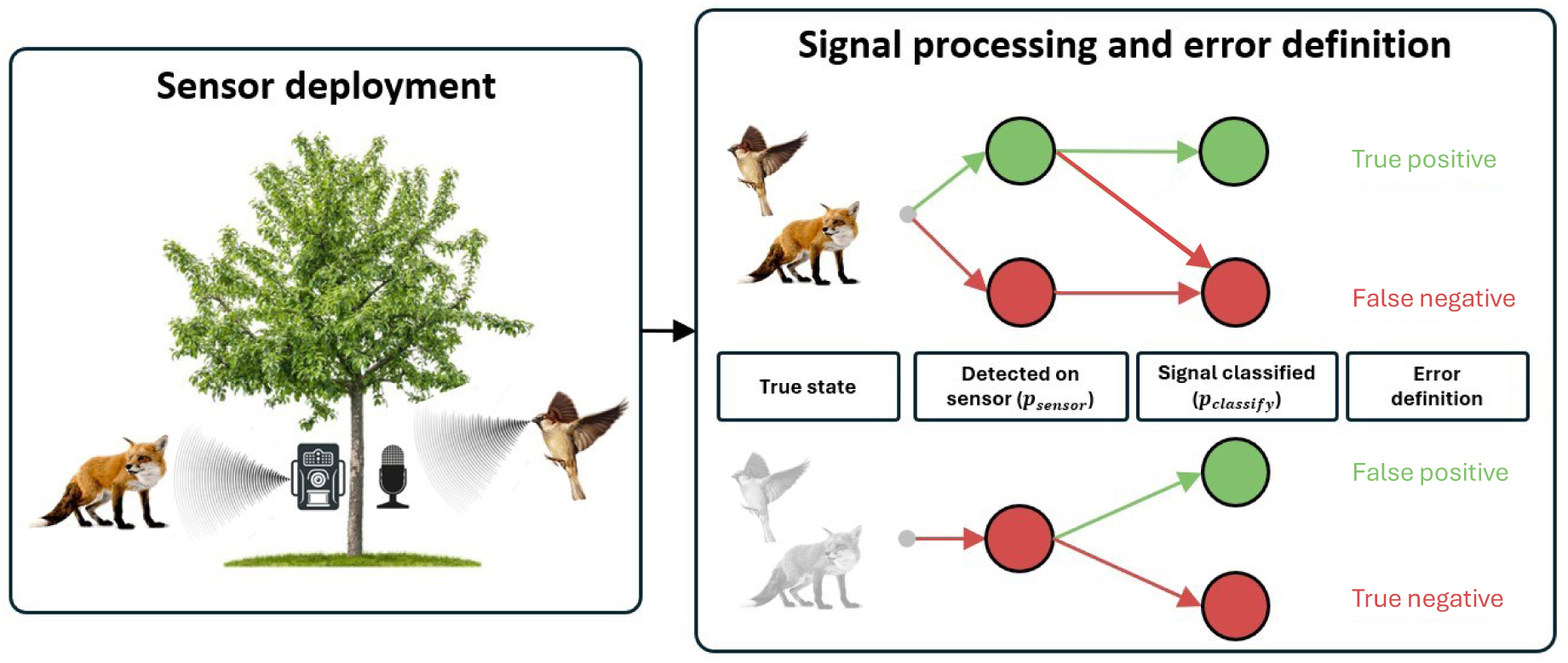
**General sources of error in sensor-based monitoring**. Sensors are deployed to monitor target species in an area (whilst we focus on autonomous recording units (ARU) and camera traps, this principle applies to any ecological sensor used to detect species presence). (1) True state: A target species (e.g., a house sparrow monitored with an ARU, or a red fox monitored with a camera trap) can be present (coloured image) or absent (greyed image) at a site. (2) Detected on sensor: If the species is present (top row), it can be detected by the sensor (with probability p_sensor_; green) or missed (with probability 1 − p_sensor_; red). If a target species is not present (bottom row), it will not be detected by the sensor (red), but other species may be detected which can later be misclassified as the target species. (3) Signal classified: Signals of the target species (green dot) can be labelled correctly (p_classify_; green arrow) or incorrectly (1-p_classify_; red arrow) as belonging to the target species. Signals not belonging to the target species (red dots) can be mislabelled as belonging to the target species (green arrow). (4) Error definition: False negatives can arise by the sensor failing to make a detection when in theory it is possible, or failure to correctly classify a signal captured by a sensor. False positives can only arise from incorrectly labelling a signal but can occur regardless of the true state.

Given the rapid uptake of automated tools alongside the emergence of new classes of statistical models, this perspective is motivated by what we perceive to be a general lack of clarity among the user community about errors associated with automated classification and modelling decisions along the sensor-data pipeline (Banner et al., 2018). As the integration of AI into ecological monitoring is in its infancy, the current lack of guidance is expected, but does result in apprehension about the accuracy, accessibility, and performance, of ecological AI. We attempt to bridge this information gap by providing clarity around the processing and modelling of AI-labelled monitoring data. We centre our perspective on occupancy modelling using the common monitoring tools of camera traps and ARUs, but note our discussions are broadly applicable to other methods (e.g. N-mixture) utilising data collected from a variety of sensors (e.g., eDNA, aerial drones). We first review the available tools that are commonly applied to acoustic and camera trap data and then discuss the mapping of data processing decisions to the appropriate statistical methods. We conclude by proposing a decision framework of suitable existing pipelines to guide ecologists through the analysis of sensor data and identify future research priorities in a rapidly developing field of research.

## Semi- and fully automated workflows for signal labelling

In what follows, we assume signals are detected, i.e., that a camera is triggered and a photo is taken or an audio recorder has recorded a sound, and that the objective is to assign the (hopefully correct) label to this ‘signal’. Manual labelling by humans will result in the assignment of a single label (e.g., a species identification) but is time intensive, resulting in bottlenecks which AI classifiers can resolve (Chalmers et al., 2023). Here we discuss options for AI-based signal labelling and the implications for their use (see Table S1 for a summary).

We define ***semi-automated workflows*** as the use of algorithms to extract general features of interest and organise large datasets into similar groups. Pre-trained neural networks applied to camera trap data label images into broad classifications. For example, the MegaDetector labels images broadly as either animal, person, or vehicle (Fennell et al., 2022). Signal recognition algorithms extract detections from lengthy audio recordings and cluster shorter sound clips by similarity (Table S1). For example, software such as KaleidoscopePro detects and clusters signals in audio clips (Guerrero et al., 2023). Even though subsequent manual labelling is required, this accurate yet broad classification improves efficiency by drastically reducing the number of signals to label (Fennell et al., 2022). Semi-automated methods may miss signals and hence increase false-negatives in the sensor detection process, but the task of general feature detection (e.g., empty, or animal, or bird call) is unlikely to produce false-positives, especially if manual validation is used.

Semi-automated clustering algorithms which attempt to increase specificity and group signals, for example from ‘animal’ to ‘species’, will be more likely to introduce false-positives. In general, the focus of what we refer to as semi-automated classification, is to detect broad rather than specific features (i.e., animal vs. specific species), removing bottlenecks but requiring a secondary labelling step.

We define ***fully automated workflows*** as the process of taking raw files as input and both detecting and labelling features of interest, thereby integrating the feature detection and labelling processes.

Full automation is computationally less efficient than the feature detection described above, but the advantage is that the output is confidence scores for specific labels (e.g., species or individual) without the need for further manual processing. Many fully automated classifiers exist (Table S1), including pre-trained classifiers for both audio recordings (e.g. BirdNET: Kahl et al., 2021) and camera trap images (e.g. Conservation AI: Chalmers et al., 2019; DeepFaune: Rigoudy et al., 2023), and deep neural network architectures that can be trained to suit user-specific purposes (e.g., images: Abadi et al., 2016, Tabak et al., 2019; audio: Lapp et al., 2023, Kahl et al., 2021).

Fully automated assignment of specific labels can introduce both false-negative and false-positive errors through misclassification. Here, it is important to make the distinction between single- and multi-species classifiers: the former assigns confidence scores to a single label of interest, whereas the latter assigns confidence scores to a suite of potential labels. For multispecies classifiers, a single signal (e.g., one image or audio clip) is assigned a non-zero confidence score for potentially many species. This introduces false-positives when a signal (i.e., an image or audio clip) has multiple non- zero labels, and false-negatives, when the true label is ignored due to higher ranking scores for incorrect labels. While the added specificity removes the need for manual labelling, fully automated algorithms are computationally more demanding and introduce more complexity in how errors emerge, and how they should be handled.

A ***two-stage ‘detect-then-label’ workflow*** is becoming increasingly common (Beery et al., 2019; Dussert et al., 2024). This process seeks to combine the efficiency of broad feature detection with the more specific task of feature labelling. This two-step process involves first applying a general feature detection algorithm (as in the semi-automated approach) and then applying the more specific classifier to the subset of data from the first step that satisfy some conditions (Rigoudy et al., 2023). The benefit is that the feature detection process can efficiently identify general features of interest (e.g, animals) thereby reducing the total number of signals to be passed to the more computationally intensive labelling process (Dussert et al., 2024). However, given the feature detection process introduces false-negative error that is typically ignored when subsets are passed to the labelling stage, this two-stage approach compounds error types – a shortcoming that appears largely overlooked in current applications.

## From signal labelling to statistical analysis

We now consider the various analytical choices available to users who have a labelled data set and wish to draw inferences about the underlying ecological process that generated the data. Traditional ecological hierarchical models were developed for binary or count data as inputs (Kéry & Royle, 2015, 2020). In the context of AI-classification where the outputs are confidence scores, the application of standard methods requires ‘thresholding’, where scores greater than or equal to a (user specified) threshold are treated as certain detections (1s), and those with less than the threshold are treated as certain non-detections (0s). Reducing classification scores to binary data ignores classification uncertainty but introduces errors with important implications for inference.

Threshold values represent a direct trade-off between precision and recall (Cole et al., 2022): high confidence thresholds will reduce false positives and increase false negatives (high precision - low recall), while low thresholds will increase false positives and reduce false negatives (low precision - high recall). This precision-recall relationship varies between different classifiers (Dussert et al., 2024) and between species classified using the same algorithm (Funosas et al., 2023; Singer et al., 2023; Whytock et al., 2021) meaning that the use of a single ‘catch-all’ threshold is unlikely to result in consistent performance between different applications and different species.

Thresholding is the most common approach when converting classification scores to data suitable for analysis with standard methods, and while bespoke workflows have been applied (e.g., Cole et al., 2022; Dussert et al., 2024; Lo Cascio et al., 2022; Symes et al., 2023; Wood et al., 2021), there is no consistent convention, and there is a clear knowledge gap around the implications of thresholding decisions. In the absence of recommendations and guidance based on simulations and experimentation, we encourage transparent reporting and justification of threshold values used and we advise against the use of a single catch-all threshold given the inherent variability in classifier specificity and labelling accuracy.

### Uncoupled classification: fitting models to binary or count data

When the labelling process is done manually, data are discrete and with good protocols, it is safe to assume that labelling errors are extremely rare (*p*_*classify*_ ≈ 1). Data can then be analysed using standard methods i.e., binary detections can be analysed using false-negative occupancy models (MacKenzie et al., 2002; Rota et al., 2016). Likewise, a widely adopted assumption is that the process of thresholding continuous score data produces binary data also readily analysed using standard methods. The most common motivation for setting high confidence thresholds is the elimination of false positive detections with a view to analysing data using models that assume only false-negative errors (Lonsinger et al., 2023; Metcalf et al., 2023; Villon et al., 2020; Whytock et al., 2021). Setting high thresholds decreases misclassification rates (Villon et al., 2020) and, as expected, improves occupancy estimates compared to non-thresholded data when using false-negative only models (Whytock et al., 2021). However, relying on high thresholds to eliminate false-positives is problematic. Conservatively high thresholds will increase false-negatives which, if detection probability is already low, can lead to bias in ecological estimates (occupancy: MacKenzie & Royle, 2005; abundance: Barker et al., 2018) and sampling inefficiencies, increasing the sampling duration required to recover reliable estimates (Cole et al., 2022). Perhaps more importantly, it is unlikely that all false-positives can be eliminated and it is well documented false-positive rates as low as 1- 3% can result in substantial biases (Clement, 2016; Royle & Link, 2006). Unlike false-negative errors, which decrease with increased temporal sampling effort, false-positive errors compound and increase with effort (McClintock et al., 2010; Miller et al., 2012; Royle & Link, 2006). This is particularly pertinent for remote sensors deployed over long periods and collect data in near-continuous time, typical of camera trap and acoustic data (Kéry & Royle, 2020).

Over the past decade, almost every class of false negative hierarchical model has been extended to account for false positive errors (e.g., occupancy: Chambert et al., 2015, Royle & Link, 2006; N-mixture: Clement et al., 2022; Doser et al., 2021), and several explicitly account for multispecies misclassification, i.e., acknowledging a false-positive detection of one species represents a false-negative in another (Augustine et al., 2023; Balantic & Donovan, 2020; Chambert, Grant, et al., 2018; Conn et al., 2014; Wright et al., 2020). These models are relatively straightforward to apply, as in many cases they have the same structure and data requirements of false negative-only models. The focus and indeed the benefit of models that account for both false-positive and false-negative errors is the ability to estimate parameters of ecological process models (e.g., occupancy or abundance) without the bias introduced by both errors (Kéry & Royle, 2020). The explicit estimation of error rates can also be of interest in certain cases. While manual labelling pipelines with robust protocols can reasonably assume perfect labelling, this is likely not the case for AI-labelled data derived from thresholding confidence scores, in which false-positives are an inherent feature. Indeed, we have observed that, although several studies report ‘consistent’ occupancy estimates when comparing models using manual vs. AI-labelled data (Gimenez et al., 2021; Lonsinger et al., 2023; Zampetti et al., 2024), there is almost always a tendency for estimates from AI-labelled data to be larger than that of manually labelled data (Box 1). This is exactly what would be expected if there were non-trivial false positive errors that were not accounted for (Cole et al., 2022).

The benefits of applying false positive models to thresholded confidence score data are two-fold. First, and most obvious, is the correction of errors that undoubtedly exist. Second, and less appreciated, is the ability to learn about error rates associated with classification that can inform and refine thresholding decisions. It is surprising that models accounting for both false-negative and false-positives are rarely used to analyse thresholded confidence score data. We argue that such models, and not false negative-only models, should be considered for such data (Box 1).

### Coupled methods: fitting models to continuous confidence score data

The desire to model uncertainties arising in each component of the observation process using automated classification has resulted in the development of ‘coupled classifiers’ for binary (Rhinehart et al., 2022) and count detections (Kéry & Royle, 2020). The coupled approach seeks to propagate uncertainty in the classification process by modelling the continuous scores directly, removing the need for thresholding. This new class of model, which is in its infancy in terms of development and application, is both conceptually and theoretically appealing. It is important to note that these models require that the full distribution of scores are retained and modelled (e.g., data cannot be truncated as many available tools do, see Rhinehart et al., 2022). Applications of the coupled approach to acoustic data have been instructive and generally encouraging but are currently restricted to single species and to scenarios unlikely to arise in ecological monitoring or research contexts (e.g., very few sites: Rhinehart et al., 2023; limited temporal sampling: Cole et al., 2022).

We are not aware of similar comparisons for camera trapping datasets. As these methods represent a significant conceptual advancement, a key focus of future research is likely to involve continued development of these models but also, we argue, better understanding of the relative merits of coupled and uncoupled methods to inform their practical use in terms of inference gains.

### Navigating the model landscape

Drawing from our experience and personal efforts to navigate the various choices available to the user community, we present a general framework to guide decision-making when modelling species detection data labelled using automated methods (Figure 2). We note that there is much research to be done to better inform these decisions and understand the importance and implications for each, and that what we provide will hopefully be updated over time.

**Figure 2.**
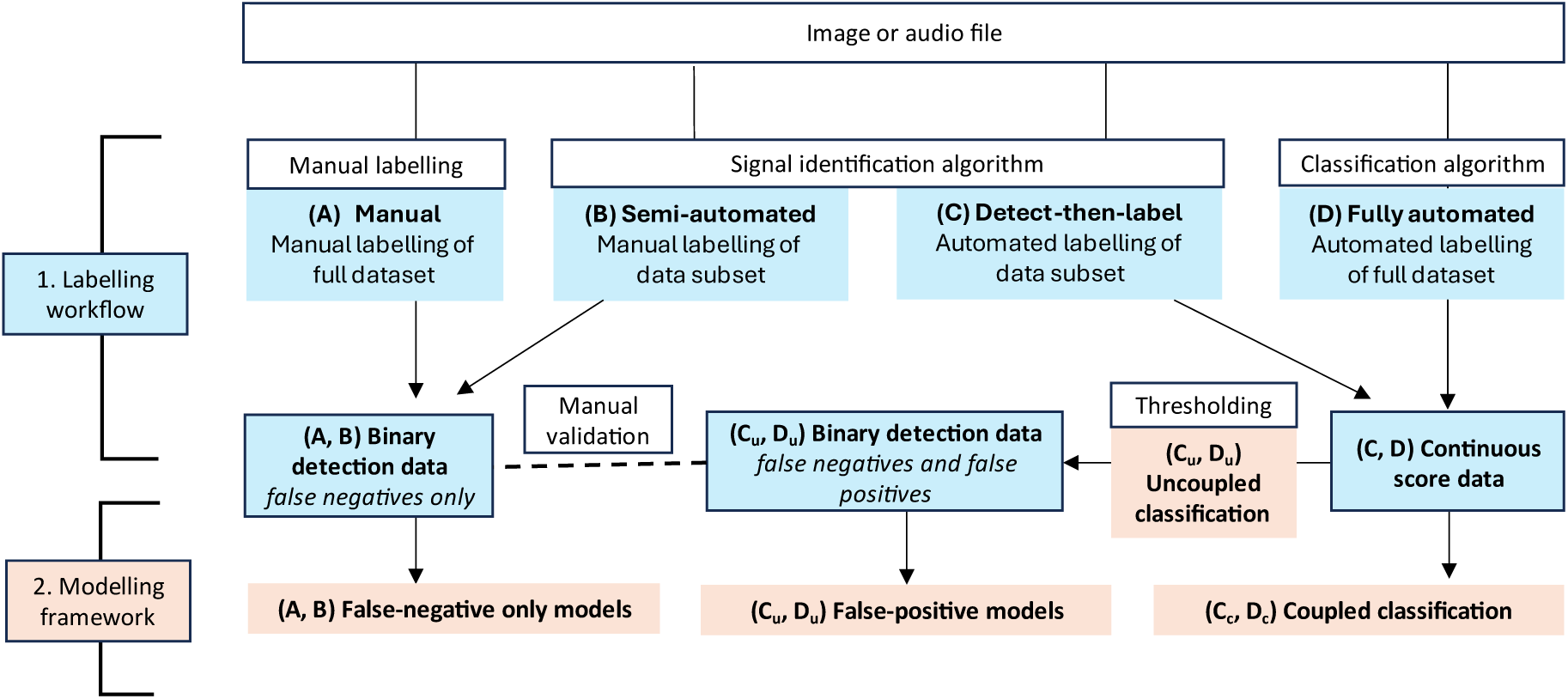
Suggested analysis pipelines for data labelled by (A) manual, (B) semi-automated, (C) detect-then-label and (D) fully automated workflows. Appropriate models for each output are mapped considering the types of errors introduced in data collection and processing. Where data are processed by (C) detect-then-label or (D) fully automated methods, the continuous score output can be analysed in coupled (denoted by subscript c) or uncoupled (denoted by subscript u) pathways.

Deciding on semi- vs. fully automated workflows should be influenced primarily by study objective (e.g., single vs. multispecies, rare vs. common species), resource requirements, and dataset size. Given human labelling costs impose practical limits on data volumes, we recommend a formal evaluation of the trade-off between human labelling cost and precision to optimise human input. Fully automated pipelines rely on training data and are often not possible if the goal is identifying a rare or data-scarce species (Guerrero et al., 2023). In this case, semi-automated methods can speed up manual labelling of relevant subsets which can eventually be used to train (ideally open access) fully automated systems for future use. Semi-automated pipelines, which typically involve some secondary manual labelling and validation of general/broad features, are likely to introduce only false-negative errors where analysis of the data using false-negative only methods is sufficient (Figure 2). Fully automated processing is less straightforward. Specifically, when thresholding, the application of false-negative only models will likely introduce (positive) bias and inflated standard errors due to the inherent and unmodelled false positives that arise through the thresholding process (Box 1). Instead, based on available methods, fully automated data should follow one of two decision branches (Figure 2). For uncoupled methods, where data is converted to binary data using a threshold, we advocate for the consideration of false-positive models as standard. If, instead, the priority is to explicitly model all sources of error (sensor and classification uncertainty), then we suggest that the confidence scores are modelled directly using coupled classifiers, but encourage readers to note the challenges discussed in the following section. We note false-negative only models can be applied to fully automated classifier output if detections with the highest confidence scores are manually validated to confirm presence or absence (e.g., Fiss et al., 2024); this is effectively a semi-automated approach. However, validating only the top scores from classifier output can elevate false-negative error and reduce the efficiency of fully automated pipelines.

Platforms such as MammalWeb, that combine the citizen science labelling capacity with automated labelling is an exemplar of a validation process that has the ability of rapid classifier validation and improvement (Hsing et al., 2022).

## Priorities for development and synthesis

While comparisons between manually- and AI-labelled data exist and are increasing in number (Gimenez et al., 2021; Lonsinger et al., 2023; Whytock et al., 2021), there is a need to formalise comparisons to motivate the uptake of specific workflows in ecology (e.g., Irvine et al., 2022). The lack of standardisation in AI-assisted monitoring pipelines has, in our opinion, contributed to a surprisingly slow uptake. Part of this challenge stems from a disconnect between the developers of AI-tools and the ecological community. Those building AI systems often focus solely on classification as the end goal with limited attention to the monitoring context, while ecologists often use AI-labelled data and proceed with traditional analyses. Bridging these disciplines is essential over the next decade as AI-technology becomes a core component of ecology in the 21^st^ century. Here, we offer some opinions on where research effort should be focussed to best capitalise on the simultaneous emergence of new computational and methodological advances.

*Thresholding confidence scores*. Optimising thresholding decisions and the benefits of doing so are currently not well understood, resulting in a lack of consistency and guidance. As there is context-dependency in the way confidence scores can be interpreted (e.g., species- or classifier-sensitivity Funosas et al., 2023; Whytock et al., 2021), thresholding based on these scores must also be context specific (Cole et al., 2022; Lo Cascio et al., 2022; Singer et al., 2023). Thresholding accuracy can be improved using auxiliary information, such as geographic and species filters (Pérez-Granados, 2023) and non-independent sequences of signals (i.e., image bursts or consecutive audio files, henceforth “sampling bursts”) which generate multiple photos or audio clips, and therefore multiple associated scores, of a single animal encounter (Singer et al., 2023; Wood et al., 2021). Filtering within these sequences can reduce errors arising from one-off high-confidence misclassifications of a singular image or clip (Wood et al., 2021), but there is uncertainty in defining sequence length, particularly for continuous audio recordings. Where ecological inference relies on modelling presence-absence across repeated extended sampling occasions (e.g., occupancy models), it is appealing to threshold data based on average scores within an extended sampling occasion (e.g., 24-hours), but the optimal method of thresholding based on signal grouping remains unclear. A better understanding of the issue of thresholding, specifically the implications for statistical inference, is a key knowledge gap.

*A base model for AI-labelled data.* A common approach for analysing continuous score data is to select a threshold, convert scores to binary detections, and apply false-negative models assuming the thresholding was sufficient to remove false-positives (Gimenez et al., 2021; Lonsinger et al., 2023; Whytock et al., 2021). We believe that using false-positive models to accommodate and quantify the classification error process should also be considered, which may reduce the influence of thresholding decisions. However, the reliability of false-positive models depends on false-positive rates being relatively low and distinguishable from false-negatives (Kéry & Royle, 2020). As false-positive models explicitly estimate the false-positive rate, there is no need to threshold data (i.e., scores > 0 can be treated as detections) unless for the purposes of speeding up model fitting (Cole et al., 2022). However, because thresholding helps reduce false-positive rates, it remains crucial to investigate and understand how thresholding choices impact the performance of false-positive models too. Considering the rapid uptake of automated methods alongside the growing use of citizen science data (Brown & Williams, 2019), false-positive errors are becoming increasingly prevalent in ecology and false-positive models in ecology should become mainstream.

Understanding the limitations of existing methods in the analysis of continuous score data (e.g., false negative models, false positive models, continuous time occupancy models) will ultimately inform future methodological development needs.

*Choosing between uncoupled and coupled analysis pipelines.* Coupled classification methods are new and as a consequence limited in their practical application (Cole et al., 2022; Kéry & Royle, 2020; Rhinehart et al., 2022). A key consideration is linking how confidence scores are generated, and the assumptions made by new and future statistical methods. While novel coupled models require all scores to effectively characterise the distributions for true and false positive detections (Rhinehart et al., 2022), many popular classifiers return scores above a certain threshold by default (e.g., conservationAI > 0.3, BirdNET > 0.01), restricting the current application of these developments.

Moreover, existing coupled methods perform poorly when score distributions are skewed. This is typical for multispecies classifiers, where every plausible species receives a score for every signal, resulting in many extremely low scores. Filtering output to keep only the top ranking species per file (e.g., Cole et al., 2022) improves this and we encourage research into defining effective filtering when applying coupled classification models to multispecies classifier output. To date, coupled classification methods have been motivated by comparisons to uncoupled false-negative approaches (Cole et al., 2022; Rhinehart et al., 2022). However, given their higher computational demands, it is key to evaluate their performance against uncoupled false-positive approaches.

*Multispecies classifiers*. Multispecies classifier output, where one signal will be assigned a confidence score for every plausible species, is typically handled by assuming the correct label is the species with the top ranking score and others may be discarded (Cole et al., 2022). However, species-specific classifier performance results in species-specific relationships between score values and precision (Funosas et al., 2023; Whytock et al., 2021). Models that take, as input, high-dimensional score vectors and model misclassification probability (e.g., Augustine, 2024) offer an important methodological development, particularly where misclassification between several species is common due to similarities in their visual or acoustic signals. These are likely to derive from similar misclassification models developed for binary or count data (Kéry & Royle, 2020). Additionally, when detections of multiple species are made in a single file (e.g., one image or audio clip), there is currently no method to determine whether high scores for two species are the result of distinct signals or misidentification of a single signal. Integrating signal recognition with labelling is likely to be beneficial in resolving this issue.

*Accessibility and co-creation*. A key focus in the ecological community has been to facilitate the pathway to opening the black-box models of machine learning. Although for developers, these considerations can improve the performance of automated algorithms, the focus of the ecological community should be redirected towards characterising and accounting for AI-associated uncertainty. Where humans are functionally capable of comprehending models using 7 ±2 rules (Sweller, 2011), it is natural to not understand the inner workings of neural networks using billions of features, and this recognition is fundamental to ensuring output is responsibly used. Our priority as an integrated user community should be (1) developing an understanding of the observation errors introduced by different automated pipelines, and (2) comparing inferential consequences of specific analytical decisions in different research contexts. This will ensure statistical frameworks to account for sensor and classifier error are fully integrated into advice on automated ecological monitoring.

## Key recommendations

The effective integration of automated methods into ecological workflows is highly beneficial but requires careful consideration of how errors arise and are handled. Manually labelled data using semi-automated workflows to maximise labelling efficiency can be analysed using false-negative only models. In comparison, fully automated workflows output confidence scores, where users can either threshold scores to construct binary data (uncoupled), or model these continuous scores directly (coupled). Thresholding decisions introduce errors that must be accounted for, and we encourage the consideration of false-positive models as well as (or instead of) false-negative only models as appropriate modelling options. As confidence scores have a species- and classifier-specific interpretation, the score-precision relationship should be quantified using validation sets (e.g., Symes et al., 2023) and if thresholds are used, they should be context-specific. Whilst evidence on the most effective method of thresholding develops, to facilitate the use of novel methods, we recommend developers of automated tools provide users with clear customisation. Successful application of false-positive models, as well as novel continuous score models, likely requires incorporation of verified files from manual validation of a subset of detections, where a degree of human identification will always remain fundamental to the successful application of automated tools (Kéry & Royle, 2020; Rhinehart et al., 2022; Royle & Link, 2006).

## Conclusions

Developing efficient and accurate analysis pipelines for ecological monitoring is essential to addressing the biodiversity crisis. Automated methods for identifying individuals and species hold great potential to transform monitoring efforts, enabling us to fully utilise the vast amount of data collected by remote sensing technologies. However, given the infancy, guidance on generating robust inference from this emerging source of species detection data remains limited; the uptake and development of relevant statistical methods has not kept pace. Motivated by our own experiences, we present a general workflow for analysing AI-labelled sensor data which maps data processing decisions to the appropriate statistical methods, and identify research avenues to facilitate the uptake of AI-assisted ecological monitoring. We emphasise that interdisciplinary co- development of workflows is far more likely to lead to innovation than shoe-horning new data streams into existing statistical methods.

---

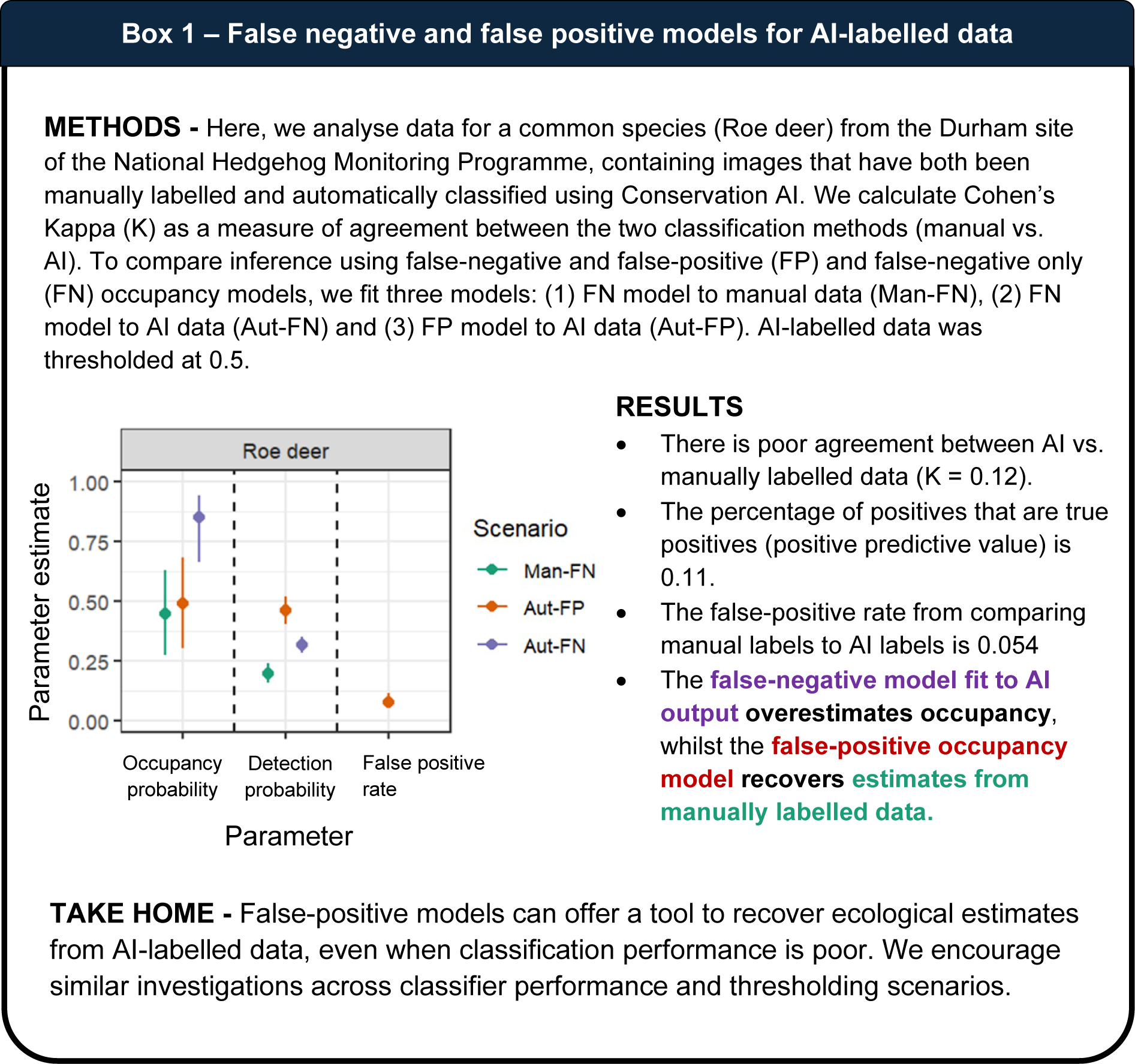

## References

1 Abadi, M., Barham, P., Chen, J., Chen, Z., Davis, A., Dean, J., Devin, M., Ghemawat, S., Irving, G., & Isard, M. (2016). {TensorFlow}: A system for {Large-Scale} machine learning. 265–283.

2. Augustine, B. (2024). Coupled Classification Occupancy. https://github.com/benaug/Coupled-Classification-Occupancy.git

3 Augustine, B. C., Koneff, M. D., Pickens, B. A., & Royle, J. A. (2023). Towards estimating marine wildlife abundance using aerial surveys and deep learning with hierarchical classifications subject to error. 10.1101/2023.02.20.529272

4 Augustine, B. C., Royle, J. A., Kelly, M. J., Satter, C. B., Alonso, R. S., Boydston, E. E., & Crooks, K. R. (2018). Spatial capture–recapture with partial identity: An application to camera traps. The Annals of Applied Statistics, 12(1). 10.1214/17-AOAS1091

5 Balantic, C. M., & Donovan, T. M. (2020). Statistical learning mitigation of false positives from template-detected data in automated acoustic wildlife monitoring. Bioacoustics, 29(3), 296–321. 10.1080/09524622.2019.1605309

6 Banner, K. M., Irvine, K. M., Rodhouse, T. J., Wright, W. J., Rodriguez, R. M., & Litt, A. R. (2018). Improving geographically extensive acoustic survey designs for modeling species occurrence with imperfect detection and misidentification. Ecology and Evolution, 8(12), 6144–6156. 10.1002/ece3.4162

7 Barker, R J , chofield, R , ink, W A , & auer, J R ( 0 8) On the reliability of N-mixture models for count data. Biometrics, 74(1), 369–377.

8. 8) Beery, S., Morris, D., & Yang, S. (2019). *Efficient Pipeline for Camera Trap Image Review* (arXiv:1907.06772). arXiv. http://arxiv.org/abs/1907.06772

9 Besson, M., Alison, J., Bjerge, K., Gorochowski, T. E., Høye, T. T., Jucker, T., Mann, H. M. R., & Clements, C. F. (2022). Towards the fully automated monitoring of ecological communities. Ecology Letters, ele.14123. 10.1111/ele.14123

10 Brown, E. D., & Williams, B. K. (2019). The potential for citizen science to produce reliable and useful information in ecology. Conservation Biology, 33(3), 561–569.

11 Burton, A. C., Neilson, E., Moreira, D., Ladle, A., Steenweg, R., Fisher, J. T., Bayne, E., & Boutin, S. (2015). REVIEW: Wildlife camera trapping: a review and recommendations for linking surveys to ecological processes. Journal of Applied Ecology, 52(3), 675–685. 10.1111/1365-2664.12432

12 Buxton, R. T., Lendrum, P. E., Crooks, K. R., & Wittemyer, G. (2018). Pairing camera traps and acoustic recorders to monitor the ecological impact of human disturbance. Global Ecology and Conservation, 16, e00493. 10.1016/j.gecco.2018.e00493

13 Caravaggi, A., Banks, P. B., Burton, A. C., Finlay, C. M. V., Haswell, P. M., Hayward, M. W., Rowcliffe, M. J., & Wood, M. D. (2017). A review of camera trapping for conservation behaviour research. Remote Sensing in Ecology and Conservation, 3(3), 109–122. 10.1002/rse2.48

14 Chalmers, C., Fergus, P., Wich, S., Longmore, S. N., Walsh, N. D., Stephens, P. A., Sutherland, C., Matthews, N., Mudde, J., & Nuseibeh, A. (2023). Removing Human Bottlenecks in Bird Classification Using Camera Trap Images and Deep Learning. Remote Sensing, 15(10), 2638. 10.3390/rs15102638

15. Chalmers, C., Fergus, P., Wich, S., & Montanez, A. C. (2019). Conservation AI: Live Stream Analysis for the Detection of Endangered Species Using Convolutional Neural Networks and Drone Technology.

16 Chambert, T., Grant, E. H. C., Miller, D. A. W., Nichols, J. D., Mulder, K. P., & Brand, A. B. (2018). Two-species occupancy modelling accounting for species misidentification and non-detection. Methods in Ecology and Evolution, 9(6), 1468–1477. 10.1111/2041-210X.12985

17 Chambert, T., Miller, D. A. W., & Nichols, J. D. (2015). Modeling false positive detections in species occurrence data under different study designs. Ecology, 96(2), 332–339. 10.1890/14-1507.1

18 Chambert, T., Waddle, J. H., Miller, D. A. W., Walls, S. C., & Nichols, J. D. (2018). A new framework for analysing automated acoustic species detection data: Occupancy estimation and optimization of recordings post-processing Methods in Ecology and Evolution, 9(3), 560–570. 10.1111/2041-210X.12910

19 Clement, M. J. (2016). Designing occupancy studies when false-positive detections occur. Methods in Ecology and Evolution, 7(12), 1538–1547. 10.1111/2041-210X.12617

20 Clement, M. J., Royle, J. A., & Mixan, R. J. (2022). Estimating occupancy from autonomous recording unit data in the presence of misclassifications and detection heterogeneity. Methods in Ecology and Evolution, 13(8), 1719–1729. 10.1111/2041-210X.13895

21 Cole, J., Michel, N., Emerson, S., & Siegel, R. (2022). Automated bird sound classifications of long-duration recordings produce occupancy model outputs similar to manually annotated data. ORNITHOLOGICAL APPLICATIONS, 124(2). 10.1093/ornithapp/duac003

22 Conn, P. B., Ver Hoef, J. M., McClintock, B. T., Moreland, E. E., London, J. M., Cameron, M. F., Dahle, S. P., & Boveng, P. L. (2014). Estimating multispecies abundance using automated detection systems: Ice-associated seals in the Bering Sea. Methods in Ecology and Evolution, 5(12), 1280–1293. 10.1111/2041-210X.12127

23 Doser, J. W., Finley, A. O., Weed, A. S., & Zipkin, E. F. (2021). Integrating automated acoustic vocalization data and point count surveys for estimation of bird abundance. Methods in Ecology and Evolution, 12(6), 1040–1049. 10.1111/2041-210X.13578

24 Dussert, G , hamaillé-Jammes, , Dray, , & iele, V ( 0 4) Being confident in confidence scores: Calibration in deep learning models for camera trap image sequences. *Remote Sensing in Ecology and Conservation*, rse2.412. 10.1002/rse2.412

25 Fennell, M., Beirne, C., & Burton, A. C. (2022). Use of object detection in camera trap image identification: Assessing a method to rapidly and accurately classify human and animal detections for research and application in recreation ecology. Global Ecology and Conservation, 35, e02104. 10.1016/j.gecco.2022.e02104

26 Fiss, C. J., Lapp, S., Cohen, J. B., Parker, H. A., Larkin, J. T., Larkin, J. L., & Kitzes, J. (2024). Performance of unmarked abundance models ith data from machine-learning classification of passive acoustic recordings. Ecosphere, 15(8), e4954.

27 Funosas, D., Barbaro, L., Schillé, L., Elger, A., Castagneyrol, B., & Cauchoix, M. (2023). *Assessing the potential of BirdNET to infer European bird communities from large-scale ecoacoustic data* [Preprint]. Ecology. 10.1101/2023.12.06.570351

28 Gimenez, O., Kervellec, M., Fanjul, J.-B., Chaine, A., Marescot, L., Bollet, Y., & Duchamp, C. (2021). Trade-off between deep learning for species identification and inference about predator-prey co-occurrence: Reproducible R workflow integrating models in computer vision and ecological statistics. arXiv Preprint arXiv:2108.11509.

29 Guerrero, M. J., Bedoya, C. L., López, J. D., Daza, J. M., & Isaza, C. (2023). Acoustic animal identification using unsupervised learning. Methods in Ecology and Evolution, 14(6), 1500– 1514. 10.1111/2041-210X.14103

30 Hsing, P., Hill, R. A., Smith, G. C., Bradley, S., Green, S. E., Kent, V. T., Mason, S. S., Rees, J., Whittingham, J , & okill, J ( 0 ) arge-scale mammal monitoring: he potential of a citizen science camera-trapping project in the United Kingdom *Ecological Solutions and Evidence*, 3(4), e12180.

31 Irvine, K. M., Banner, K. M., Stratton, C., Ford, W. M., & Reichert, B. E. (2022). Statistical assessment on determining local presence of rare bat species. Ecosphere, 13(6), e4142. 10.1002/ecs2.4142

32 Kahl, S., Wood, C. M., Eibl, M., & Klinck, H. (2021). BirdNET: A deep learning solution for avian diversity monitoring. Ecological Informatics, 61, 101236. 10.1016/j.ecoinf.2021.101236

33 ) Kéry, M., & Royle, J. A. (2015). Applied hierarchical modeling in ecology: Volume 1: Prelude and static models. Elsevier Science.

34 ) Kéry, M., & Royle, J. A. (2020). Applied hierarchical modeling in ecology: Analysis of distribution, abundance and species richness in R and BUGS: Volume 2: Dynamic and advanced models. Chapter 7. Academic Press.

35 Lapp, S., Rhinehart, T., Freeland-Haynes, L., Khilnani, J., Syunkova, A., & Kitzes, J. (2023). OpenSoundscape: An open-source bioacoustics analysis package for Python. Methods in Ecology and Evolution, 14(9), 2321–2328. 10.1111/2041-210X.14196

36 Leorna, S., & Brinkman, T. (2022). Human vs. machine: Detecting wildlife in camera trap images. Ecological Informatics, 72, 101876. 10.1016/j.ecoinf.2022.101876

37 Lo Cascio, A., Kasel, S., & Ford, G. (2022). A new method employing species-specific thresholding identifies acoustically overlapping bats. Ecosphere, 13(11), e4278. 10.1002/ecs2.4278

38 Lonsinger, R. C., Dart, M. M., Larsen, R. T., & Knight, R. N. (2023). Efficacy of machine learning image classification for automated occupancy-based monitoring *Remote Sensing in Ecology and Conservation*, rse2.356. 10.1002/rse2.356

39 MacKenzie, D. I., Nichols, J. D., Lachman, G. B., Droege, S., Andrew Royle, J., & Langtimm, C. A. (2002). ESTIMATING SITE OCCUPANCY RATES WHEN DETECTION PROBABILITIES ARE LESS THAN ONE. Ecology, 83(8), 2248–2255. 10.1890/0012-9658(2002)083[2248:ESORWD]2.0.CO;2

40 MacKenzie, D. I., & Royle, J. A. (2005). Designing occupancy studies: General advice and allocating survey effort. Journal of Applied Ecology, 42(6), 1105–1114.

41 McClintock, B. T., Bailey, L. L., Pollock, K. H., & Simons, T. R. (2010). Unmodeled observation error induces bias when inferring patterns and dynamics of species occurrence via aural detections. Ecology, 91(8), 2446–2454.

42. Metcalf, O., Abrahams, C., Ashington, B., Baker, E., Bradfer-Lawrence, T., Browning, E., Carruthers-Jones, J., Darby, J., Dick, J., & Eldridge, A. (2023). *Good practice guidelines for long-term ecoacoustic monitoring in the UK*.

43 Miller, D. A., Weir, L. A., McClintock, B. T., Grant, E. H. C., Bailey, L. L., & Simons, T. R. (2012). Experimental investigation of false positive errors in auditory species occurrence surveys. Ecological Applications, 22(5), 1665–1674.

44 Palencia, P., Barroso, P., Vicente, J., Hofmeester, T. R., Ferreres, J., & Acevedo, P. (2022). Random encounter model is a reliable method for estimating population density of multiple species using camera traps. Remote Sensing in Ecology and Conservation, 8(5), 670–682. 10.1002/rse2.269

45 Pérez-Granados, C. (2023). BirdNET: applications, performance, pitfalls and future opportunities. Ibis, 165(3), 1068–1075. 10.1111/ibi.13193

46 Rhinehart, A , urek, D , & Kitzes, J ( 0 ) A continuous-score occupancy model that incorporates uncertain machine learning output from autonomous biodiversity surveys. Methods in Ecology and Evolution, 13(8), 1778–1789. 10.1111/2041-210X.13905

47 Rigoudy, N., Dussert, G., Benyoub, A., Besnard, A., Birck, C., Boyer, J., Bollet, Y., Bunz, Y., Caussimont, G., Chetouane, E., Carriburu, J. C., Cornette, P., Delestrade, A., De Backer, N., Dispan, L., Le Barh, M., Duhayer, J., Elder, J.-F., Fanjul, J.-B , … hamaillé-Jammes, S. (2023). The DeepFaune initiative: A collaborative effort towards the automatic identification of European fauna in camera trap images. European Journal of Wildlife Research, 69(6), 113. 10.1007/s10344-023-01742-7

48 Rota, C., Wikle, C., Kays, R., Forrester, T., McShea, W., Parsons, A., & Millspaugh, J. (2016). A two-species occupancy model accommodating simultaneous spatial and interspecific dependence. ECOLOGY, 97(1), 48–53. 10.1890/15-1193.1

49 Royle, J. A., & Link, W. A. (2006). GENERALIZED SITE OCCUPANCY MODELS ALLOWING FOR FALSE POSITIVE AND FALSE NEGATIVE ERRORS. Ecology, 87(4), 835–841. 10.1890/0012-9658(2006)87[835:GSOMAF]2.0.CO;2

50 Singer, D., Hagge, J., Kamp, J., Hondong, H., & Schuldt, A. (2023). Aggregated time-series features boost species-specific differentiation of true and false positives in passive acoustic monitoring of bird assemblages. 10.13140/RG.2.2.10556.82568

51 ) Sweller, J. (2011). Cognitive load theory. In Psychology of learning and motivation (Vol. 55, pp. 37–76). Elsevier.

52 Symes, L., Sugai, L. S., Gottesman, B., Pitzrick, M., & Wood, C. (2023). Acoustic analysis with BirdNET and (almost) no coding: Practical instructions. 10.5281/ZENODO.8357176

53 Tabak, M. A., Norouzzadeh, M. S., Wolfson, D. W., Sweeney, S. J., Vercauteren, K. C., Snow, N. P., Halseth, J. M., Di Salvo, P. A., Lewis, J. S., White, M. D., Teton, B., Beasley, J. C., Schlichting, P. E., Boughton, R. K., Wight, B., Newkirk, E. S., Ivan, J. S., Odell, E. A., Brook, R. K , … iller, (2019) Machine learning to classify animal species in camera trap images: Applications in ecology. Methods in Ecology and Evolution, 10(4), 585–590. 10.1111/2041-210X.13120

54 Villon, S., Mouillot, D., Chaumont, M., Subsol, G., Claverie, T., & Villéger, S. (2020). A new method to control error rates in automated species identification with deep learning algorithms. Scientific Reports, 10(1), 10972. 10.1038/s41598-020-67573-7

55 Whytock, R , Ś ieże ski, J , Z erts, J A , Bara- łupski, , Koumba Pambo, A F , Rogala, , Bahaa-el-din, , Boekee, K , Brittain, , ardoso, A W , Henschel, P , ehmann, D , Momboua, B., Kiebou Opepa, C., Orbell, C., Pitman, R. T., Robinson, H. S., & Abernethy, K. A. (2021). Robust ecological analysis of camera trap data labelled by a machine learning model. Methods in Ecology and Evolution, 12(6), 1080–1092. 10.1111/2041-210X.13576

56 Wood, C. M., Kahl, S., Chaon, P., Peery, M. Z., & Klinck, H. (2021). Survey coverage, recording duration and community composition affect observed species richness in passive acoustic surveys. Methods in Ecology and Evolution, 12(5), 885–896. 10.1111/2041-210X.13571

57 Wright, W. J., Irvine, K. M., Almberg, E. S., & Litt, A. R. (2020). Modelling misclassification in multi-species acoustic data when estimating occupancy and relative activity. Methods in Ecology and Evolution, 11(1), 71–81. 10.1111/2041-210X.13315

58 Zampetti, A., Mirante, D., Palencia, P., & Santini, L. (2024). Towards an automated protocol for wildlife density estimation using camera-traps. bioRxiv, 2024–08.

